# Stability of feature selection utilizing Graph Convolutional Neural Network and Layer-wise Relevance Propagation

**DOI:** 10.1101/2021.12.26.474194

**Authors:** Hryhorii Chereda, Andreas Leha, Tim Beißbarth

**Affiliations:** Medical Bioinformatics, University Medical Center Göttingen, Germany; Medical Statistics, University Medical Center Göttingen, Germany; Campus-Institute Data Science (CIDAS), University of Göttingen, Göttingen, Germany

**Keywords:** gene expression data, explainable AI, personalized medicine, precision medicine, classification of cancer, deep learning, prior knowledge, molecular networks

## Abstract

High-throughput technologies are increasingly important in discovering prognostic molecular signatures and identifying novel drug targets. Molecular signatures can be obtained as a subset of features that are important for the decisions of a Machine Learning (ML) method applied to high-dimensional gene expression data. However, feature selection is inherently unstable in this case. Several studies have identified gene sets that provide predictive success for patient prognosis, but these sets usually have only a few genes in common. The stability of feature selection (and reproducibility of identified gene sets) can be improved by including information on molecular networks in ML methods. Graph Convolutional Neural Network (GCNN) is a contemporary deep learning approach applicable to gene expression data structured by a prior knowledge molecular network. Layer-wise Relevance Propagation (LRP) and SHapley Additive exPlanations (SHAP) are techniques to explain individual decisions of deep learning models. We used both GCNN+LRP and GCNN+SHAP techniques to explain GCNNs and to construct feature sets that are relevant to models by aggregating their individual explanations. We also applied more classical ML-based feature selection approaches and analyzed the stability, impact on the classification performance, and interpretability of selected feature sets.

**Availability:** https://gitlab.gwdg.de/UKEBpublic/graph-lrp

## 1 Introduction

Microarray and especially high-throughput technologies have become commonly used tools for genome-wide gene-expression profiling. Gene expression patterns elucidate the molecular mechanisms of such heterogeneous disease as breast cancer (Sørlie, 2007). As a result, large amounts of data produced by high-throughput sequencing are utilized to identify predictive gene signatures and discover individual biomarkers in cancer prognosis (Perera, Leha, and Beissbarth, 2019). To predict cancer outcomes, we can use ML to classify patients, and identify relevant genes by selecting features.

However, an ML model, trained on a high-dimensional gene expression dataset (the number of genes is much higher than the number of patients), has to deal with the “curse of dimensionality”, which leads to overfitting (Porzelius et al., 2011) and consequent instability in feature selection. Furthermore, the correlations between single genes and the clinical endpoint fluctuate strongly when measured over different subsets of patients (Ein-Dor et al., 2005) affecting the lists of candidates for predictive biomarkers. From the biomedical standpoint, it is crucial to guarantee the reproducibility of the given feature selection methods when finding candidate sets of biomarkers (Lee et al., 2013). Thus, the challenge is to find a subset of important features that are (i) stable (reproducible), (ii) biologically interpretable, (iii) contain the most information about the clinical outcome.

The stability (reproducibility) of a feature selection algorithm is essentially the robustness of the algorithm’s feature preferences: the feature selection is unstable when small changes in the training data lead to large changes in the chosen feature subsets (Nogueira, Sechidis, and Brown, 2018). The quantification of stability can be performed by providing different samples from the same training data and measuring the changes among chosen feature subsets.

Incorporation of prior knowledge of molecular networks (e.g. pathways) into an ML algorithm improves stability (Johannes et al., 2010) and biological interpretability (Kong and Yu, 2018, “Introduction” section of) of selected gene signatures. Molecular networks represent molecular processes in a given biological system and are widely used by biologists to interpret the results of a statistical analysis (Porzelius et al., 2011). Molecular networks can be used to approximate the interactions between features (genes). Genes connected in a close proximity within molecular network should have similar expression profiles (Johannes et al., 2010) and should not be treated independently.

The Graph Convolutonal Neural Network (GCNN) (Defferrard, Bresson, and Vandergheynst, 2016) have been successfully applied to classify patients utilizing their gene expression profiles structured by a prior knowledge protein-protein interaction (PPI) network (Rhee, Seo, and Kim, 2018; Chereda et al., 2019; Ramirez et al., 2020; Pfeifer et al., 2023). Patient-specific predictions were further explained (Chereda et al., 2021) by adapting the Layer-wise Relevance Propagation (LRP) explanation method (Bach et al., 2015) to the graph convolutions layers of the GCNN method.

Here, we aggregate individual explanations delivered by LRP and SHAP (Lundberg and Lee, 2017) applied to a GCNN model to construct a feature set corresponding to a whole dataset. We estimate the stability of feature selection performed by the GCNN+LRP and GCNN+SHAP approaches w.r.t. different training samples. Comparing GCNN+LRP, GCNN+SHAP and other more classical ML-based feature selection techniques we analyzed the stability, impact on the classification performance, and interpretability of feature sets of varying size.

## 2 Materials and Methods

### 2.1 Protein-Protein Interaction Network

The gene expression data was structured with the Human Protein Reference Database (HPRD) protein-protein interaction (PPI) network (Keshava Prasad et al., 2009). It contains protein-protein interaction information based on yeast two-hybrid analysis, in vitro and in vivo methods. The set of binary interactions between pairs of proteins in the HPRD PPI network represented as an undirected graph. The graph has several connected components.

### 2.2 Breast Cancer Data

We utilized RNA-seq based gene expression dataset of human breast cancer patient samples. Each patient in this dataset is labeled with a molecular breast cancer subtype. Batch normalized expression (Illumina HiSeq_RNASeqV2) and clinical data are provided by The Cancer Genome Atlas (TCGA) and were downloaded from (*cBioPortal TCGA-BRCA* PanCancer data 2018). The expression data comprise the collection of 20531 genes and 1082 samples. After mapping sample IDs to clinical data (containing subtype labels) we ended up with 981 samples in breast cancer, corresponding to five subtypes: luminal A (499 samples), luminal B (197 samples), basal-like (171 samples), HER2-enriched (78 samples) and normal-like (36 samples).

The gene expression data was normalized utilizing the gene length corrected trimmed mean of M-values (GeTMM) method (Smid et al., 2018). It allows for inter- and intra-sample analyses with the same normalized data set. After that we applied *log*_2_(*x*+1) transform to reduce the scale. The expression data were mapped to vertices of PPI resulting in 8469 genes in the main connected component. Since GCNN requires an underlying graph to be connected, the rest of the genes were mostly singletons, thus not included in analyses.

### 2.3 Methods for feature selection

We use GCNN, Multi-Layer Perceptron (MLP), and Random Forest (RF) (Breiman, 2001) for predictive modelling. SHAP and LRP explanation methods were applied to the same GCNN and MLP models to generate individual explanations. These explanation were aggregated according to the the approach of (Marcílio and Eler, 2020) to deliver model-wide feature sets. The methods are described in the following subsections.

#### 2.3.1 Predictive modelling with GCNN, MLP, and RF

The GCNN method (Defferrard, Bresson, and Vandergheynst, 2016) was applied to the breast cancer dataset (“2.2 Breast Cancer Data”). The HRPD PPI network (“2.1 Protein-Protein Interaction Network”) represents connections between genes and was used to structure gene expression data. The topology of the HRPD PPI network is used by GCNN and is the same for each patient. For a single patient, each vertex of the HPRD PPI network has a corresponding gene expression value as an attribute. A gene expression profile of a single patient creates a graph signal and GCNN performs graph signal classification.

MLP and RF were applied on the same dataset performing classification of patients, but without utilizing network topologies.

#### 2.3.2 LRP and SHAP explanation methods

The LRP and SHAP methods were developed for delivering data point specific explanations. We apply those methods to MLP and GCNN models.

LRP computes a relevance value for each feature (gene) of an individual data point (cancer patient) by backpropagating the full relevance (output label) through the layers of a neural network to its input. To compute relevance values we utilized the *z*^+^ propagation rule (Montavon et al., 2017) that was applied to each layer of GCNN and MLP.

The SHAP method estimates Shapley values, a term coming from cooperative game theory. According to Molnar, 2019, the game theory setup behind Shapley values is the following: The “game” is the prediction task for a single data point. The “payout” is the difference between the actual prediction for this data point and the average prediction for all instances. The “players” are the feature values of the data point that collaborate to receive the “payout” (predict a certain value). Shapley values indicate how to fairly distribute the “payout” among the features. SHAP’s DeepExplainer approach was applied to our Keras (Chollet, 2015) implementation of GCNN (Chereda, 2022).

A single Shapley value or relevance value computed by LRP represents how important a particular feature value is to a decision made by a classifier for a specific data point.

#### 2.3.3 Aggregating individual explanations from LRP and SHAP to deliver a feature set

A single explanation can be represented as a vector of relevances or a vector of Shapley values estimates respectively. If gene expression data

*X* ∈ *R*^*n×m*^ is given as an input to be explained, then the rows of the matrix *E* = (*e*_*ij*_) ∈ *R*^*n×m*^ are transposed vectors of single explanations. *n* is the number of data points and *m* is the number of features. Let *C* be the set of classes, and |*C*| is the number of classes. Marcílio and Eler, 2020 suggest an approach to score features by aggregating SHAP values generated for each class of the dataset. We apply the same approach to aggregate relevence values computed by LRP. The aggregating approach is the following (Marcílio and Eler, 2020)

1. Generate explanations {*E*_*c*_: *c ∈ C}* for each class and take element-wise absolute values 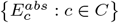. The latter set of matrices we combine into tensor

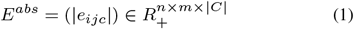

for explicitness of further notations.
2. For each matrix 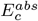 one computes mean values column-wise (feature-wise). The result is transposed vectors

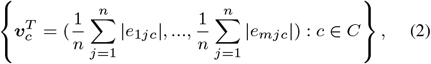

which we combine into matrix 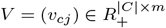.
3. Let vector ***q*** be a representation of importance scores. The importance scores are obtained by summing vectors *v*_*c*_, e.g.

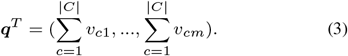

The features then are sorted according to the importance scores. Fixing *t* as a number of features to select, one can construct a set of most important features *s*, |*s*| = *t*

Within 10-fold cross validation, GCNN or MLP was trained on training data and the explanations are generated by LRP or SHAP on test data. LRP and SHAP methods were applied to exactly the same models.

#### 2.3.4 Feature selection with SHAP and RF

Important features of a RF model were selected using two approaches: standard scoring of features on the basis of mean decrease in Gini impurity and application of SHAP’s TreeExplainer to RF. The SHAPley values were generated on test sets within 10-fold cross validation and the features were selected in the same as way as described above.

### 2.4 Measuring the stability of feature selection

The input of a feature selection procedure is the data set 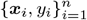 where *n* is number of data points, each ***x***_*i*_ is an *m*-dimensional feature vector, and *y*_*i*_ is the associated label. Feature selection identifies a feature subset *s* of dimensionality *k < m* (Nogueira, Sechidis, and Brown, 2018). The subset *s* is selected to convey the most relevant information about the label *y* for the whole dataset 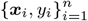. The output of a feature selection approach is either a scoring on the features, a ranking of the features, or a subset of the features. Further in this paper, we do not consider the scoring information about features selected and treat them as a set.

The input dataset of a feature selection technique is a finite sample that is created by a generating distribution. In the case of varying samples, the selected feature subset may vary as well. The variation of the feature subset is the stability that we aim to measure.

A typical approach to measure stability is to produce *M* subsamples of the dataset at hand, to apply a feature selection approach to each one of them, and then to measure the variability in the *M* feature sets obtained (Nogueira, Sechidis, and Brown, 2018). Let *Z* = {*s*_1_, …, *s*_*M*_ } be a collection of feature sets. Let *ϕ*(*s*_*i*_, *s*_*j*_) be a symmetric function taking two feature sets as input and returning their similarity value and let 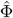 be a function taking *Z* as input and returning a stability value. Nogueira, Sechidis, and Brown, 2018 provide a good overview over stability measuring techniques. We utilize a similarity based approach, so that 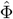 can be defined as the average pairwise similarity between the *M* (*M* − 1)*/*2 possible pairs of feature sets in *Z*:

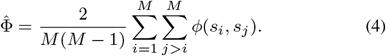

As an easily interpretable pairwise similarity function, we use the Jaccard distance:

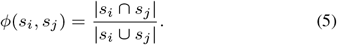

One of the techniques to generate M subsamples from a dataset is bootstrap. Another approach is random subsampling (Wald, Khoshgoftaar, and Dittman, 2012). We utilize 10-fold cross validation to generate data subsamples, therefore *M* = 10. SHAP and LRP methods were applied to explain test data, and explanations were aggregated into 10 selected feature sets. We estimate the stability using equation (4) for different sizes |*s*_*i*_| varying from 100 to 1000 with a step 100.

### 2.5 Evaluating the impact of features prioritized by LRP and SHAP on classification performance

We adapt a “pixel-flipping” procedure (Montavon, Samek, and Müller, 2018; Bach et al., 2015) to quantify the impact of selected features on classifier’s performance by perturbing them. Within 10-fold cross-validation, we perturb features {*s*_1_, …, *s*_10_} of test sets by assigning their values zero, which would mean the absence of detected gene expression of particular genes in a gene expression profile of a patient in a test set. Then we apply ML models to perturbed test data and record an average of the predictive performance over 10 folds. Increasing the size of perturbed selected feature sets decreases classifier’s performance. We compare the drop of the classification importance across MLP+LRP, MLP+SHAP, GCNN+LRP, GCNN+SHAP, RF+SHAP, RF approaches by varying the size of a feature set |*s*_*i*_| from 100 to 1000 with a step 100.

### 2.6 Evaluating connectivity of the selected feature sets

We estimate the connectivity of selected feature (gene) sets as a measure of interpretability in the context of the prior knowledge HRPD PPI network. We build a PPI subnetwork, in which nodes are selected genes, and edges are from a PPI network, connecting those genes. Further, we count the number of connected components of the resulting PPI subnetwork. Lower number of connected components indicates higher connectivity, and thus better interpretability within a prior knowledge HPRD PPI network. The number of connected components is averaged over 10 feature sets obtained within 10-fold cross-validation. We repeat measurements varying the size of the feature set |*s*_*i*_| from 100 to 1000 with a step 100.

While only GCNN uses prior knowledge HRPD PPI network, we have also quantified the connectivity of feature sets delivered by MLP and RF methods without prior knowledge for comparison.

## 3 Results

### 3.1 Stability of feature selection

The stability metrics demonstrate (Figure 1) that the feature selection using GCNN+LRP (violet) is substantially more stable than using RF, MLP, or GCNN+SHAP. Classical feature selection with RF (blue) provides high stability as well and it almost the same as the stability of the RF+SHAP approach (orange). GCNN+SHAP (brown) shows slightly higher stability than that of MLP+SHAP (red) and MLP+LRP (green), which is the lowest. Interestingly, the stability slightly drops for RF with increased feature size, compared to other methods.

**Fig. 1.**
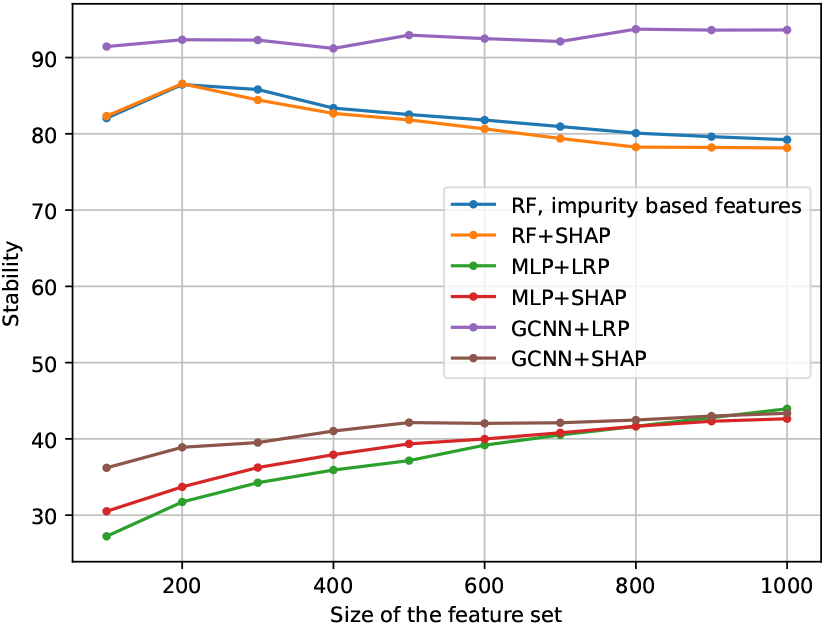
The stability of feature selection. Higher is better.

### 3.2 The impact of selected features on classification performance

We first evaluated the ML methods using a 10-fold cross validation. Table 1 shows the results. GCNN and MLP outperformed Random Forest. Later, we proceeded with the computational experiments evaluating the impact of selected features on classifier’s performance (Figure 2). For smaller feature sizes, the largest drop in classification performance demonstrate features selected by RF (blue and orange) and GCNN+SHAP (brown) approaches. GCNN+LRP (violet) features show rather moderate drop in performance compared to GCNN+SHAP (brown) features. As for the MLP model, MLP+SHAP features (red) also show higher impact on performance than MLP+LRP features (green). Note, the pairs of explanation methods were always applied to the same models.

**Table 1.**
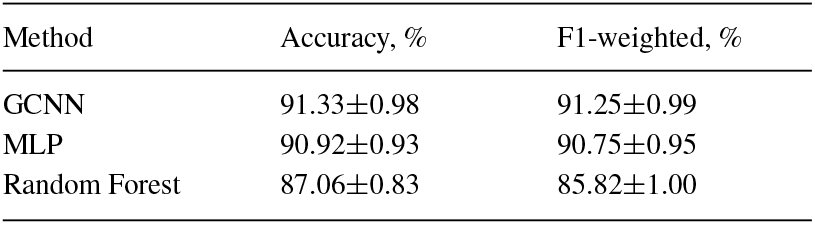
Performance of ML models predicting breast cancer subtypes.

**Fig. 2.**
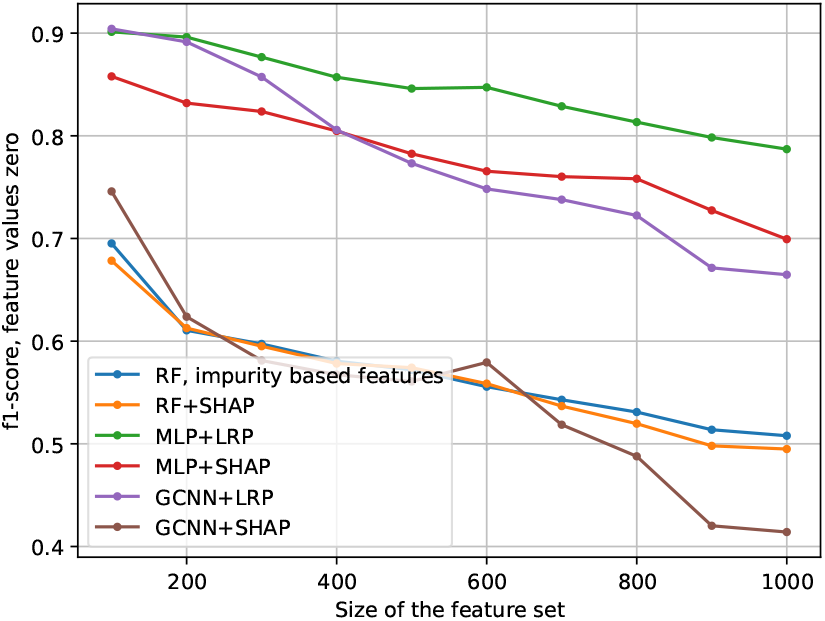
The impact of selected features on classifier’s performance. The selected features are perturbed in test sets within 10-fold cross validation and averaged F1-weighted classification scores are recorded. Lower is better.

### 3.3 The connectivity of selected features

Feature sets selected by GCNN+LRP have the lowest number of connected components (highest connectivity) within HPRD PPI network compared to that of feature sets selected by other methods (Figure 3). For example, for the feature size |*s*_*i*_| = 200 the average number of connected components is equal to 11 for GCNN+LRP, while other methods provide *s*_*i*_ with the average number of connected components higher than 122. The connectivity of GCNN+SHAP features resembles that of MLP+SHAP features. Classical feature selection with RF provides almost the same connectivity of features as using RF+SHAP. The number of connected components of the MLP+LRP features grows slower than that of features selected by other methods except for GCNN+LRP.

**Fig. 3.**
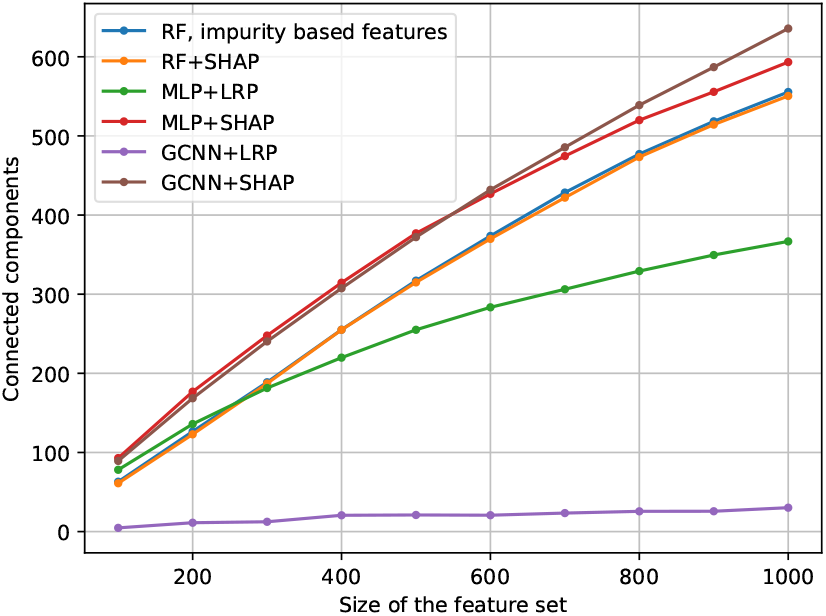
The connectivity is represented by the average (within 10-fold cross validation) number of connected components in a HPRD PPI subnetwork constructed from a selected feature set. Lower is better (more interpetable in a context of HPRD PPI network).

## 4 Discussion

The focus of our paper is to holistically investigate the stability, impact on the classification performance, and interpretability of feature sets obtained by aggregating individual explanations from LRP and SHAP methods. We contrast the GCNN+LRP and GCNN+SHAP approaches that use HPRD PPI network as prior knowledge with other techniques such as MLP+LRP, MLP+SHAP, RF+SHAP, and RF (standard featuere scoring) that do not use any prior knowledge.

GCNN+LRP leads in stability and connectivity of features, while GCNN+SHAP produces features that are more impactful for classification but less interpretable in terms of prior knowledge. RF also generates impactful features with high stability. One possible reason for GCNN+SHAP’s behavior is that SHAP’s DeepExplainer approach, which approximates SHAP values, assumes the input features are independent (Lundberg and Lee, 2017). However, DeepExplainer still uses a GCNN model that leverages network neighborhoods. To better understand this phenomenon, we would need to conduct more experiments using other explanation methods, such as integrated gradients (Sundararajan, Taly, and Yan, 2017) or different LRP rules (see (Montavon et al., 2019, Appendix A of), Kohlbrenner et al., 2020), but we leave this for future work.

The MLP+SHAP and MLP+LRP features provide less steep line than GCNN+LRP and GCNN+SHAP in the context of the impact on classification performance (Figure 2). It can be explained by the assumption that the MLP model in this case relies on different patterns learned from data, since patterns learned by GCNN are biased by the PPI networks. Furthermore, one has to take into consideration the non-triviality of estimating the impact of features on performance by perturbing them. For example, perturbed features might lead to artifacts, and it might be unclear if those artifacts originated from data, or a misbehaving model, or misbehaving explanation method (Sundararajan, Taly, and Yan, 2017). To minimize this effect, we perturbed features by making them equal zero and used the same models to which LRP and SHAP were applied. One could also consider an adaptation to graph domain a more advanced empirical approach described in (Samek et al., 2017).

There are other feature selection approaches that could be considered in the same context. Ahn et al., 2018 suggest an empirical approach for analyzing gene’s contribution to a particular output of a deep neural network model. This approach could be also potentially applicable to GCNN. Kong and Yu, 2018 present a graph-embedded feed-forward network for feature selection. The model they propose is essentially an MLP where the first hidden embedding layer has its weights pruned by an adjacency matrix of the utilized molecular network. For feature selection they also developed a feature ranking method, which require input gene expression data to be standardized. In our case, the input data is not standardized because we use GeTMM method (Smid et al., 2018) for data normalization. It allows for inter- and intra-sample analyses. Standardization treats genes independently distorting gene expression patterns formed in the same neighborhood of a molecular network which in turn would affect the interpretability of features (Chereda et al., 2021). The GNNSubNet approach from (Pfeifer, Saranti, and Holzinger, 2022) allows for feature selection utilizing a modified version of GNNExplainer (Ying et al., 2019) applied to the Graph Isomorphism Network GIN (Xu et al., 2018). GIN shows good performance in a multimodal setup (Pfeifer, Saranti, and Holzinger, 2022) however it does not converge well while utilizing only gene expression modality (see also Pfeifer, Saranti, and Holzinger, 2022, Table 2). Nonetheless, adapting a modified version of GNNExplainer to our Keras (Chollet, 2015) implementation of GCNN (Chereda, 2022) is a development for future work.

While here we are concentrated on aggregating data point-specific explanations into a general scoring for features, there is no “gold standard” for performing general feature selection using explanation methods. The GCNN+LRP approach can deliver explanations as patient-specific subnetworks (Chereda et al., 2021) and it might be promising to work with subnetworks, rather than aggregating SHAP or relevance values. Authors in (Lapuschkin et al., 2019) perform spectral relevance analysis by clustering individual explanations, and in a graph domain one could cluster patient-specific subnetworks (Chereda et al., 2021) to deliver a model-wide subnetwork consisting of important features.

In addition to graph neural network-based approaches, there are methods that use ensembles of trees and leverage information from molecular networks: (Dutkowski and Ideker, 2011) and (Pfeifer et al., 2022). While this is a fascinating topic to cover, it is out of scope for this paper.

## 5 Conclusion

We evaluated different feature selection techniques for identifying prognostic gene sets: GCNN+LRP and GCNN+SHAP (using HPRD PPI network as prior knowledge), MLP+LRP, MLP+SHAP, RF+SHAP, and RF (using no prior knowledge). We examined how the size of the feature sets affects their stability, classification performance, and interpretability. RF+SHAP and RF (standard scoring) produced similar and relatively stable and impactful features. GCNN+LRP yielded the most stable and interpretable gene sets among all used in this paper ML methods, making it more suitable for biomarker discovery. MLP+LRP was the least reliable for feature selection as it had the lowest stability. GCNN+SHAP generated features that were less stable and interpretable than GCNN+LRP, but had more influence on the model’s performance. The SHAP method, applied to both MLP and GCNN, is useful when the features need to be more relevant to the model’s decisions.

## Acknowledgements

We would like to acknowledge Johannes Söding for fruitful discussions. T.B. is a member of the Göttingen Campus Institute Data Science.

